# Spatiotemporal Models Reveal Dynamic Growth Patterns in U.S. West Coast Groundfish

**DOI:** 10.1101/2025.05.19.653526

**Authors:** Andrea N. Odell, Kristin N. Marshall, Eric J. Ward, Kelli F. Johnson, Marissa L. Baskett

## Abstract

Variability in somatic growth of marine fish affects their reproductive potential and survival, and therefore, the productivity of a population. Understanding how growth might vary among species with similar traits can improve predictions of population status and responses to environmental change. We used geostatistical models to characterize spatiotemporal variability in growth rate and body condition, two interrelated traits associated with somatic growth, of nine commercially important U.S. West Coast groundfish species with a range of life-history traits and depth distributions. Our models uncover spatiotemporal variability in growth rate and body condition in all nine groundfish species with limited trends shared among species with similar traits, suggesting a greater influence from niche partitioning acting on local scales. Such interspecific differences in growth rate and body condition responses also occurred at regional scales, with some species exhibiting positive responses while others declined. These findings reveal the dynamic nature of somatic growth among groundfish species and provide insight into potential mechanisms of its variability that could be considered within climate-enhanced assessments of population status for marine fish.

## Introduction

In an ecosystem-based approach to fisheries management that accounts for variable ocean conditions in space and time (Botsford *et al*., 1997), there is growing recognition that variation in demographic traits influences the productivity and dynamics of marine fish populations. Identifying patterns and drivers of variability in demographic traits can improve estimates of productivity and set expectations following environmental change (Stawitz *et al*., 2019; Correa *et al*., 2021). Somatic growth is one demographic trait that varies over space and through time in marine fishes (Stawitz *et al*., 2015; Gertseva *et al*., 2017). It contributes to reproductive potential, sexual maturity, and survival (Peters, 1986) and governs the productivity and density of a population (Lorenzen and Enberg, 2002). In some cases, the effect of growth variation on population outcomes is more than that of recruitment variation (Stawitz and Essington, 2018) but recruitment variation is more often incorporated into stock assessments (Maunder and Thorson, 2019). As such, incorporating variable growth into population models has the potential to reduce bias and increase both accuracy in estimates of population status (Punt *et al*., 2020; Correa *et al*., 2021) and predictability of population response to changing environmental conditions (Pinsky and Mantua, 2014).

Somatic growth is regulated by the dynamic relationship between length, weight, and age and can be characterized using a variety of metrics. Two common metrics are length-at-age and body condition, and their variability can reflect the environmental, ecological, and fishing influences that a species experiences at a given location and time. Local environmental conditions can directly affect food availability and physiological processes, which might either promote growth when conditions are favorable or limit growth when conditions are not (Champion *et al*., 2020; Cominassi *et al*., 2020). Ecological interactions within a community (e.g., strength of competition for food resources) can shift in response to direct environmental influences and affect growth indirectly (Clark *et al*., 2018). Moreover, fishing can induce artificial selection toward smaller length-at-age and earlier maturation (Heino *et al*., 2015) which can alter population structure (Cheung *et al*., 2013) and increase population sensitivity to environmental perturbations (Ratner and Lande, 2001). The relative importance of such environmental, ecological, and fishing influences can fluctuate across space and time, potentially creating a complex mosaic of variability in somatic growth across a species’ distribution (Thorson, 2015; Gertseva *et al*., 2017) and lifespan (Stawitz *et al*., 2015), which can be difficult to disentangle.

Variability in responses of somatic growth to environmental change might depend on a species’ life history and spatial distribution. Long-lived species typically exhibit slower ecological responses to environmental change compared to short-lived species (Thresher *et al*., 2007), likely attributed to their weaker ability to compensate for environmental change through rapid demographic response (Perry *et al*., 2005). However, such environmental change does not occur uniformly across space. For example, temperatures increase most rapidly at the surface and these rates decline with depth, suggesting that warming might play a weaker role in somatic growth variability of deeper-dwelling species. According to metabolic theory, fish are expected to exhibit faster growth rates and larger size-at-age of young individuals as temperatures warm, which are traits associated with greater productivity that might buffer populations against the negative effects of fishing (Atkinson, 1994). Experimental and observational studies support these patterns predicted by metabolic theory (Huss *et al*., 2019; Lindmark *et al*., 2022b) but evidence also supports both slower or variable growth with warming temperatures when life histories are faster (e.g., faster growth, earlier maturity, higher natural mortality; Wang *et al*., 2020) and when prey availability decreases (Rovellini *et al*., 2024; Liu *et al*., *In review*).

One class of commonly-harvested species with demonstrated spatiotemporal variability in growth are groundfish; examples include Atlantic cod (Frater *et al*., 2019), Pacific cod (Correa *et al*., 2020), Antarctic Toothfish (Webber and Thorson, 2016), and rockfishes along the U.S. West Coast (Harvey *et al*., 2011). These species occupy a broad distribution of depths with varied life-history traits and have supported commercial and recreational fisheries for more than a century (Miller *et al*., 2014). While life history and depth distribution are theorized to indicate a species’ growth response to environmental change, growth in groundfish species might be further influenced by trophic niche partitioning where prey resources are used differently among species with large overlaps in their spatial distributions (Garrison and Link, 2000). Such niche partitioning might obscure the expected response in growth shared among sympatric species with similar life history and depth distributions, complicating our understanding of its variability in groundfish.

Along the U.S. West Coast, the California Current is a highly productive ecosystem that supports many marine fisheries including groundfish species that are relatively well-sampled with previous studies documenting variability in their growth. Spatial variability in asymptotic length and growth rate were observed along a discretized latitudinal gradient in eight groundfish species using von Bertalanffy length-at-age models (Gertseva *et al*., 2017). These and other groundfish species also show evidence of temporal variability in size-at-age at the regional scale (Stawitz *et al*., 2015; Thorson and Minte-Vera, 2016) and body condition at the local scale (Thorson, 2015). Growth rate and body condition are interrelated, and limited evidence suggests a tradeoff might exist between them for some species (Karametsidis *et al*., 2023).

Following these analyses of groundfish growth rate and body condition, the U.S. West Coast experienced an unprecedented marine heatwave that persisted between 2014 and 2016, and the frequency of marine heatwaves has since increased (Jacox *et al*., 2022). Such extreme environmental anomalies can reduce food availability (Gomes *et al*., 2024) and affect somatic growth (Smith *et al*., 2023). Therefore, the U.S. West Coast groundfish present a promising system to investigate the drivers of spatiotemporal variation in multiple traits and at multiple spatial scales for a range of life histories and habitats.

In this paper, we develop a model-based framework for estimating relative changes in somatic growth, measured as growth rate and body condition, and apply this novel framework to expand our understanding of spatiotemporal variation in somatic growth in U.S. West Coast groundfish species. In contrast to previous studies that apply biogeographical or jurisdictional breaks to estimate growth in discrete spatial blocks (Gertseva *et al*., 2017), we model space as a continuous process to better capture the nature of similarities shared between neighboring locations compared to more-distant locations (Anderson *et al*., 2024). With this geostatistical model, we summarize our findings on the range and patterns of spatiotemporal variation in growth rate and body condition for nine groundfish species. We compare and contrast patterns between species with different life-history traits and depth distributions. Finally, we discuss potential mechanisms that might be attributed to these observed patterns.

## Methods

### Description of data

The Northwest Fisheries Science Center (NWFSC) has conducted an annual West Coast Groundfish Bottom Trawl Survey (WCGBTS; Keller *et al*., 2017) since 2003 to obtain biological, distributional, and abundance information of over 90 species included in the Pacific Fishery Management Council’s Groundfish Fishery Management Plan. The survey operates along the continental shelf and upper slope (from 55 to 1280 m depth) between Cape Flattery, Washington and the U.S. (N 48° 22’)–Mexico border (N 32° 34’). Typically, two coastwide passes are conducted by four industry-chartered vessels using an Aberdeen style trawl net (standard four-panel, single bridle, with a small mesh codend liner), sampling approximately 650 sites annually selected via a stratified random sampling design (Keller et al. 2017). The first pass occurs from mid-May to late July, and the second pass occurs from mid-August to late October. Because of the global pandemic, surveys were not conducted in 2020.

Trends in growth rate or body condition can be difficult to quantify for species that are rare or not well-sampled by the survey; therefore, we removed species that were observed fewer than 500 times across all years. From the data that remained, we omitted observations that did not record sex, weight, length, or age. For the majority of species sampled in the WCGBTS, ages are estimated from otoliths. Aging is both time intensive and expensive – as a result, otoliths are not processed from all species in all years and are primarily processed just prior to stock assessments. As there is also a delay in processing otoliths, age information might not be complete in the most recent years despite the availability of otoliths. Consequently, some species have an incomplete time series of age estimates. After this filtering process, nine species were selected representing a range of different life histories (Table 1). Though age and size information exist for both males and females, we focused only on female individuals because many marine fish allocate energy differently between sexes causing sexual dimorphism. This resulted in 54,147 observations of females with paired length, weight, and age measurements across the nine species. Each species had at least 150 samples for each year that samples were available and the spatial coverage of sampling aligned with their known distribution. The length of the time series varied among species ranging between 6 and 19 years of samples. See Supplementary Materials for more details.

**Table 1.** Analyzed species and their designation of depth distribution, lifespan, and maturation, and estimates of maximum size measured using total length obtained from FishBase. * indicates female-specific maximum size. For specific values of maximum age and age at 50% maturity, see Table S2.

| Species | Depth distribution | Lifespan | Maturation | Maximum size (L<sub>inf</sub>) |
| --- | --- | --- | --- | --- |
| Aurora rockfish | Slope | Long-lived | Late-maturation | 41 cm |
| Dover sole | Shelf–slope | Long-lived | Early-maturation | 76 cm |
| Pacific sanddab | Shelf–slope | Short-lived | Early-maturation | 41 cm |
| Petrable sole | Shelf | Short-lived | Early-maturation | 70 cm* |
| Darkblotched rockfish | Shelf–slope | Long-lived | Late-maturation | 58 cm |
| Splitnose rockfish | Slope | Long-lived | Late-maturation | 46 cm |
| Lingcod | Shelf | Short-lived | Early-maturation | 152 cm |
| Pacific hake | Shelf–slope | Short-lived | Early-maturation | 105 cm* |
| Sablefish | Shelf–slope | Long-lived | Early-maturation | 120 cm |

We selected dover sole (*Microstomus pacificus*), Pacific sanddab (*Citharichthys sordidus*), petrale sole (*Eopsetta jordani*), aurora rockfish (*Sebastes aurora*), darkblotched rockfish (*Sebastes crameri*), splitnose rockfish (*Sebastes diploproa*), lingcod (*Ophiodon elongatus*), Pacific hake (*Merluccius productus*), and sablefish (*Anoplopoma fimbria*). These species inhabit a range of depth distributions, over the continental shelf (shelf species), deeper along the slope (slope species), or more broadly distributed across the shelf and slope (shelf–slope species). In addition, they capture a wide range of life histories, long-lived species with late maturation, short-lived species with early maturation, and long-lived species with early maturation. We grouped species into life history and depth-distribution categories (Gertseva et al., 2017; Thorson, 2015; Table 1) to draw comparisons among species.

We obtained information on female maximum (total) length for each of the nine species from FishBase, a global information system on fishes (Boettiger *et al*., 2012). We accessed this database using the *rfishbase* package (v 4.1.2) in R (R Core Team, 2023).

### Spatiotemporal models

#### Growth rate model

We modeled growth rate as a hierarchical process with spatial, temporal, and spatiotemporal random effects. We estimated growth rates for each species by fitting a linearized version of the von Bertalanffy growth function (VBGF) to length and age data from the WCGBTS (Figure 1). The basic form of the VBGF models describe the expected length at age, *L*(*a*), given an estimated asymptotic length *L*_∞_, Brody growth coefficient *k*(yr^-1^), and age when average length is zero *a*_0_ following the form 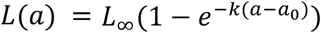. This model can be extended to include a time-varying *k*_*t*_, where 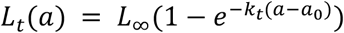. From here, we can linearize the time-varying VBGF by rearranging and applying log transformations to both sides, such that

**Figure 1.**
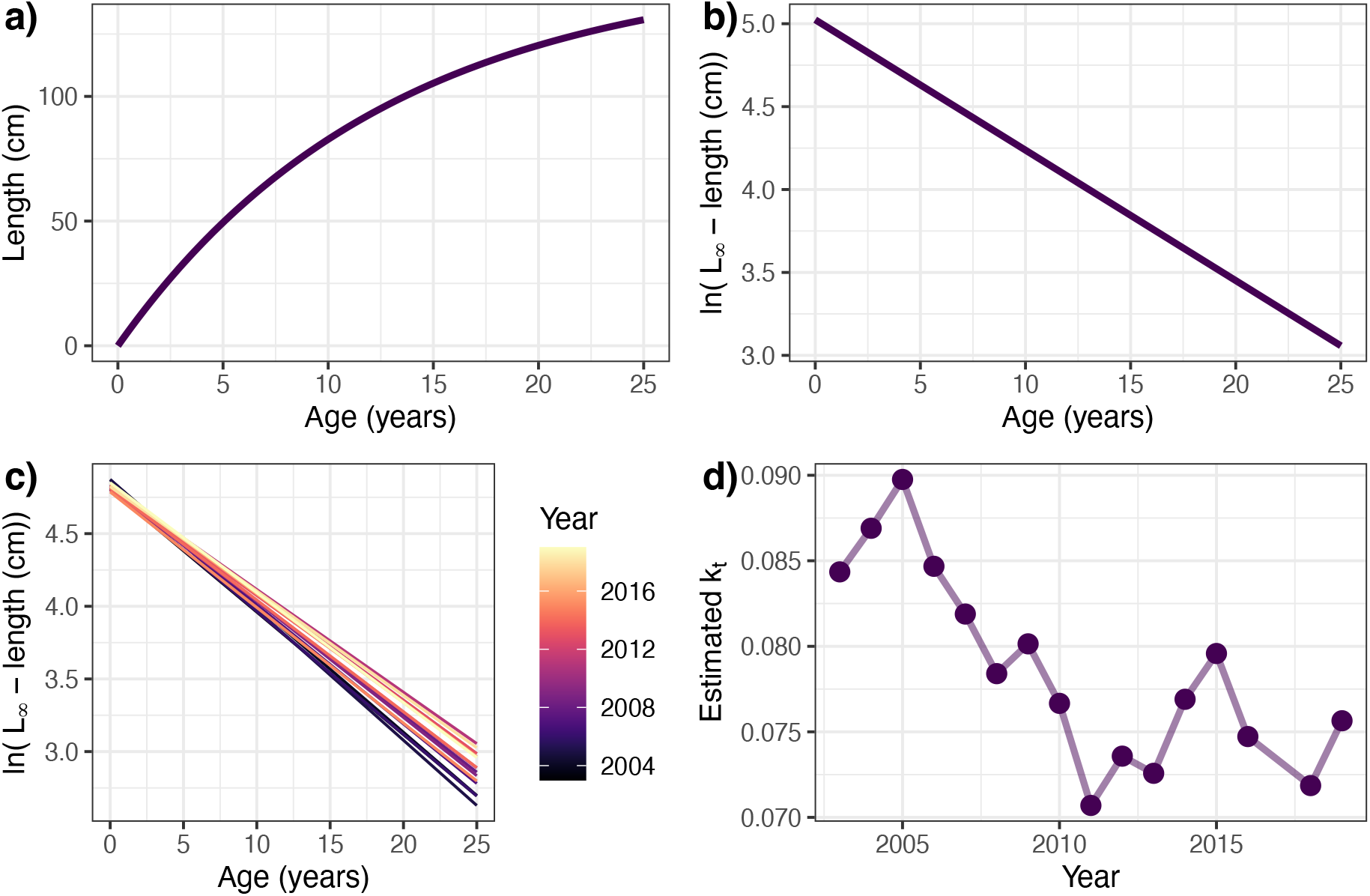
Conceptual illustration of the time-varying linearized von Bertalanffy growth function, using biological data from the northern subpopulation of lingcod (*Ophiodon elongatus*). The estimated von Bertalanffy growth function (a) is linearized by applying log transformations to both sides of the equation (b). A unique function can be estimated for each year (or location) as indicated by a continuous color scale by fitting time-varying random effects for the slope and intercept(c). The slope of the function for each year represents the estimated growth coefficient, *k*_*t*_, as it varies over time (d). Here, we used a simplified version of the estimation model for illustrative purposes, which only captures temporal variability and ignores spatial locations.

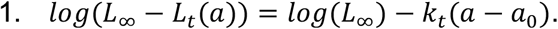

In this linear form, *log*(*L*_∞_− *L*_*t*_(*a*)) becomes the response variable, *log*(*L*_∞_) can be described using the intercept, and −*k*_*t*_ can be described using the slope (Figure 1b). Because our data represent a relatively brief snapshot in time, we assumed *L*_∞_ to be constant and greater than the maximum observed size (Table 1). We then calculated log2*L*_∞_− *L*_*t*_(*a*)) for each observation which was used to fit the model. We estimated variability between years as random effects on the slope and intercept. We account for differences due to survey timing using sampling day of year as data are collected over the span of five months within a survey year. We assume that *a*_0_ represents age 0.

While the choice of *L*_∞_ or *a*_0_ directly affects the absolute estimates of growth rate *k*, it does so in a consistent manner such that estimated growth coefficients from this model can be used to identify relative trends in time or space.

To extend equation 1 to account for spatial and spatiotemporal variation in the mean growth rate in each year, we used the Stochastic Partial Differential Equation (SPDE) approximation to Gaussian Markov Random Fields (GMRF; Lindgren *et al*., 2011) and estimated spatial random effects as well as spatially-varying random effects on age *a* for each year *t*. In doing so, we can account for location and year-specific conditions that may influence each age uniquely. This is one way to account for movement in mobile species within the VBGF model such that their size-at-age reflects a cumulation of their growth environments throughout their lifespan (He and Bence, 2007). We generated a triangulated mesh within the spatial domain of the WCGBTS using a cutoff distance of 30 km (points separated by smaller values being assigned to the same knot or vertex; Rue *et al*., 2009) resulting in a mesh with 192 knots.

We fit a spatiotemporal model for each species by calculating and using the log-transformed difference between *L*_∞_ and observed length-at-age *L*_*s,t*_(*a*) in year *t* and location *s* as the response variable, 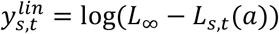, and assumed normally distributed residuals. By estimating 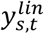 with a linear model, we derived *k* by taking the negative of the estimated slope (Equation 1). In this setting, our spatiotemporal VBGM can be expressed as a geostatistical generalized linear mixed model (GLMM):

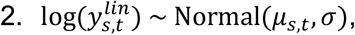

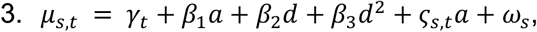

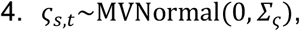

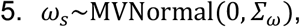

where *µ*_*s,t*_ represents the mean log-transformed difference between *L*_∞_ and observed length-at-age *L*_*s,t*_(*a*) at location *s* and time *t*, and *σ* represents the residual standard deviation. We explained variation in *µ*_*s,t*_ with the following covariates: *γ*_*t*_ represents independent global intercepts estimated for each year (as a factor), *a* represents age, *d* represents day of year (centered and scaled) modeled as a quadratic function for its flexibility, and *β*_*i*_ where *i* = 1,2,3 represents fixed effect coefficients. We include *ω*_*s*_ and *ς*_*s,t*_ which, respectively, are standard variables used to represent the spatial field with covariance matrix *Σ*_*ω*_ and the spatially-varying coefficients on age for time *t* which varies with location *s* with covariance matrix Σ_*ς*_.

All spatially-varying random effects, *ω*_*s*_ and *ς*_*s,t*_, are modeled as a GMRF with covariance matrices constrained by an anisotropic Matérn function that defined how the rate of spatial covariance decays with distance given a parameter that controls smoothness, ν, fixed at 1 (Lindgren *et al*., 2011) and an estimated parameter that represents the spatial decorrelation rate, *κ*. For the spatially varying coefficients, we assumed the same variance across years − this allows for year-specific spatial effects but forces the magnitude of spatial variability in growth to remain static over time. Together, the coefficients on age (*β*_1_, *ς*_*s,t*_) can be interpreted as controlling growth in a similar way to *k*_*t*_ in the non-spatial VBGF described above. This spatiotemporal approach to estimating growth coefficients borrows information across years to estimate an apparent year-specific growth coefficient. An alternative model structure could be to fit spatiotemporal models to length data for each age and subtract them to obtain a more direct estimate of growth rate, however sample sizes in our current data set preclude that approach.

#### Body condition model

We estimated female body condition across space and time using Le Cren’s relative condition factor 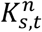 (Le Cren, 1951), where the superscript *n* distinguishes this body condition metric from Fulton’s condition factor *K*. This factor characterizes deviations from an expected weight given observed length as an indication of body condition and is calculated as the ratio between the observed weight of an individual in location *s* and time *t, w*_*s,t*_, and a predicted weight, 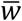, given an observed length. Therefore, 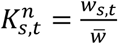 such that an individual with 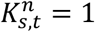 exhibits a body condition that is average across all years and locations. This approach overcomes some of the limitations of other condition metrics such as the assumption of isometric growth used by Fulton’s *K* (Lloret *et al*., 2013). We estimated a predicted weight, 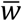, given an observed length, *l*, following the length−weight relationship 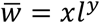 where we estimated the parameters *x* and *y* using a simple non-spatiotemporal model (see Supplementary Materials). We then divided all observed weights, *w*_*s,t*_, by 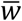 to calculate the Le Cren’s relative condition factor, 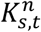. By standardizing to 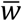 generated from a simple non-spatiotemporal model, all deviations across space and time (e.g., seasonality or food availability) will be captured in the derived 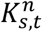 values.

For each species, we used log-transformed 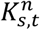 as the response variable and assumed student-t distributed residuals. We continued with the same GLMM framework used for growth rate and expressed our spatiotemporal body condition models as:

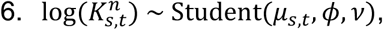

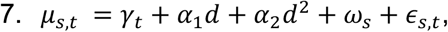

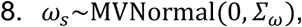

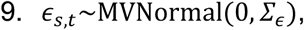

where *µ*_*s,t*_ represents the mean condition factor, *ϕ* represents the scale parameter, and ν represents the available degrees of freedom. We included covariates *γ*_*t*_ and *d* similar to the growth rate model (Equation 3) and *α*_*i*_ where *i* = 1,2 are fixed effect coefficients. We included *ω*_*s*_ and *ϵ*_*s,t*_, which represent spatial and spatiotemporal random fields. We used the same GMRF approach described above to model spatial and spatiotemporal variability with covariance matrices *Σ*_*ω*_ and *Σ*_*ϵ*_, respectively, and modeled independent spatiotemporal random fields for each year. We centered and scaled year *t* in *γ*_*t*_ and *ϵ*_*s,t*_ in this model to improve model convergence.

#### Biomass model

When generating annual indices of growth and body condition, we applied a biomass-weighting factor to account for spatial heterogeneity in a species’ distribution when estimating indices at the regional-scale. The weighting factor provided greater emphasis on local estimates where biomass of a species is high compared to where biomass is low. We created a biomass-weighting factor for each location and year by estimating biomass using aggregated catch and effort data from the WCGBTS. We used catch-per-unit-effort (kg per km^2^) as the response variable and assumed the residuals followed a Tweedie distribution with a log-link function to better handle zero-inflated data (Tweedie, 1984; Shono, 2008). We continued with the same GLMM framework and expressed our spatiotemporal biomass models as:

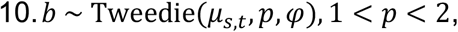

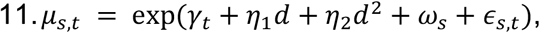

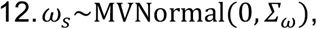

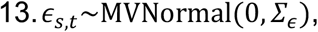

where *µ*_*s,t*_ represents the mean biomass, *p* represents the power parameter, and *φ* represents the dispersion parameter. We include independent global intercepts *γ*_*t*_, fixed coefficients *η*_1_ and *η*_2_ for day of year *d* modeled as a quadratic effect, and *ω*_*s*_ and *ϵ*_*s,t*_ which represent spatial and spatiotemporal random effects, respectively. We assume *ω*_*s*_ and *ϵ*_*s,t*_ are drawn from GMRFs with covariance matrices *Σ*_*ω*_ and *Σ*_*ϵ*_, respectively. We considered alternative weighting schemes such as weighting by numbers of individuals instead of biomass and chose biomass weighting because this method integrates biomass across ages and sizes in a particular location, while weighting by numbers would likely overrepresent younger ages that are more frequently observed for most species.

### Parameter estimation

We fit models using Template Model Builder (TMB; Kristensen *et al*., 2016), which integrates over random effects using the Laplace approximation. TMB calculates standard errors on all estimated quantities using a generalized delta-method. We fit all models using the *sdmTMB* package (Anderson *et al*., 2024) in R version 4.3.2 (R Core Team), a flexible interface that links R-INLA and TMB under a single package. We assessed convergence by verifying two criteria, (1) that fitted models resulted in a Hessian matrix that was positive definite and (2) that the absolute log likelihood gradient with respect to fixed effects was <0.001.

### Spatial predictions and index generation

We used each of the fitted models for growth rate, relative body condition, and biomass (for index-weighting) to predict values onto a regularized grid and, for each species, to evaluate spatiotemporal variation and generate model-based indices. We used a 4×4 km grid for spatial predictions, similar to the survey design grid (Keller et al. 2017) available from the *nwfscSurvey* package (Wetzel *et al*., 2024). We used all unique values of covariates to make predictions, except for day of year which we predicted to July 1st of each calendar year as it is the approximate median date of survey sampling in the data we used.

We derived growth rates in each spatial cell using predictions from the growth model where the slope of the linearized VBGF is −*k*_*s,t*_. When visualizing spatial variation, we normalized *k*_*s,t*_ so the values scaled between 0 and 1. This normalization allowed us to compare spatial patterns of variability between species but not the range of variability. In this case, normalized *k*_*s,t*_ values that are close to 0 represent locations where individuals exhibit the slowest growth rates among that species, whereas values closer to 1 represent locations where individuals exhibit the fastest growth rates of the species.

To generate model-based growth indices, we weighted *k*_*s,t*_ by the proportion of the total biomass at time *t, B*, predicted in that grid cell. This weighting factor, 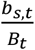, where *b*_*s,t*_ represented the predicted biomass in location *s* and time *t*, is included to account for spatiotemporal heterogeneity in biomass, giving more weight to growth estimates in cells with higher biomass. We then calculated an annual biomass-weighted growth index by summing up biomass-weighted *k*_*s,t*_ across the entire spatial domain each year. An exception was made for lingcod because of their strong genetic differentiation along the coast, for which we generated a northern and a southern biomass-weighted growth index split atN 40° 10’ (Longo *et al*., 2020). We generated metrics of uncertainty around growth by passing the fitted sdmTMB model to Stan using Rstan (Stan Development Team 2024a and 2024b) and running 4 parallel Markov chain Monte Carlo (MCMC) chains (1000 warmup iterations, 1000 sampling iterations).

Because of computational challenges in estimating uncertainty around derived quantities of interest, we fixed most parameters at point estimates and focused on propagating uncertainty via MCMC samples of the spatially varying effects of age, *ς*_*s,t*_, because these represent spatiotemporal variation in our model.

Unlike the growth rate model, 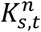 was the predicted output from its model and did not need further derivation. We weighted 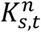 by *b*_*s,t*_ as described above to generate an annual index of biomass-weighted body condition,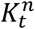. Again, the spatial domain of lingcod was split into northern and southern subpopulations at N 40° 10’ before generating the biomass-weighted body condition. As 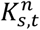 was not a derived quantity, we generated uncertainty around the predicted condition using the same MCMC sampling procedure used for growth. All code and data to reproduce our analysis is on our Github repository, https://github.com/AndreaNOdell/groundfish_growth.

## Results

### Range of variation in growth rate

Groundfish species along the U.S. West Coast exhibited interspecific differences in how widely growth rate and body condition varied across time and space. Consistent patterns among species with similar life history traits did not emerge for the groundfish species included in this study (Figure 2). All short-lived species exhibited greater spatiotemporal variation compared to spatial variation, though the difference in Pacific sanddab was small (Figure 2a). Dover sole and sablefish, both long-lived early maturing species, exhibited slightly greater spatial variation than spatiotemporal variation. While aurora rockfish and splitnose rockfish exhibited relatively equal amounts of spatial and spatiotemporal variation in growth rate, darkblotched rockfish exhibited much more spatiotemporal than spatial variation despite also being a long-lived late maturing species.

**Figure 2.**
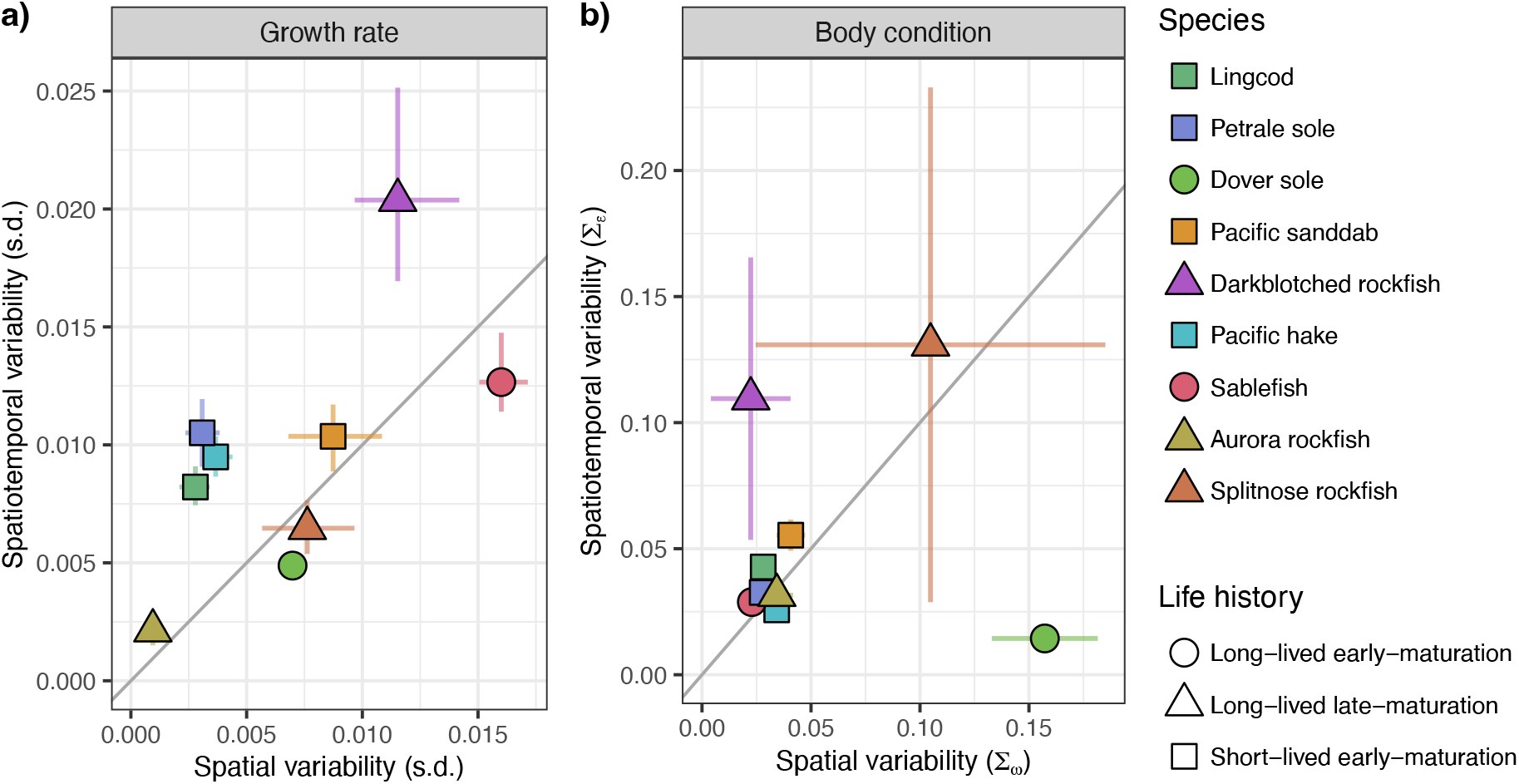
Spatial and spatiotemporal variability in growth rate *k* (a) and body condition *K*^*n*^ (b) for species (colors) that are long-lived and early maturing (circles), long-lived and late maturing (triangles), and short-lived and early maturing (squares). Variability in *k*is measured as the standard deviation (s.d.) calculated from estimates of *k* and variability in *K*^*n*^ is measured as the standard deviation of the spatial and spatiotemporal fields estimated by the body condition model, *Σ*_*ω*_ and *Σ*_s_, respectively. Error bars indicate uncertainty as confidence intervals for growth rate and as the estimated standard error of *Σ*_*ω*_ and *Σ*_s_ for body condition. The 1:1 line indicates whether spatiotemporal variability (above) or spatial variability (below) is more influential.

For body condition, most species exhibited relatively comparable amounts of spatial and spatiotemporal variation (Figure 2b). However, two long-lived species displayed markedly stronger variation in body condition along one axis: dover sole showed greater spatial variation while darkblotched rockfish exhibited greater spatiotemporal variation. Maturation schedules differentiate these responses, where dover sole exhibits earlier maturation compared to darkblotched rockfish. While we did not observe consistent patterns in growth rate or body condition associated with depth distribution, we cannot make any inferences because only two species were attributed to the shelf and to the slope (Figure S1).

### Spatial variation across species

Specific locations in which growth rate was favorable between species did not appear to be consistent among species with similar life history or depth distribution (Figure 3). Shelf species did not show similar patterns overall, lingcod displayed strong patchiness along the coast, whereas petrale sole exhibited pronounced hotspots of fast growth rates localized off the northwest tip of Washington state and at the outlet of San Francisco Bay. Slope species, which were both long-lived with late maturation (aurora and splitnose rockfish), demonstrated a latitudinal gradient with faster growth rates observed in the northern half of their range. Among the more widely distributed shelf– slope species, sablefish and dover sole shared similar regions of fast growth, with their fastest growth rates occurring along the Washington and Oregon coastline and decreasing further offshore. These species also share similar life-history characteristics, as they are both long lived with early maturation. Pacific hake and darkblotched rockfish, both shelf–slope species with contrasting life histories, exhibited strong patchiness in growth rates, akin to lingcod, with patches of their slowest growth occurring in the northern part of their range. The absence of patchiness in southern California for darkblotched rockfish was likely due to their minimal presence in that region. Pacific sanddab exhibited the most prominent latitudinal gradient, with their highest growth rates observed in the central part of their range and minimal variation with depth.

**Figure 3.**
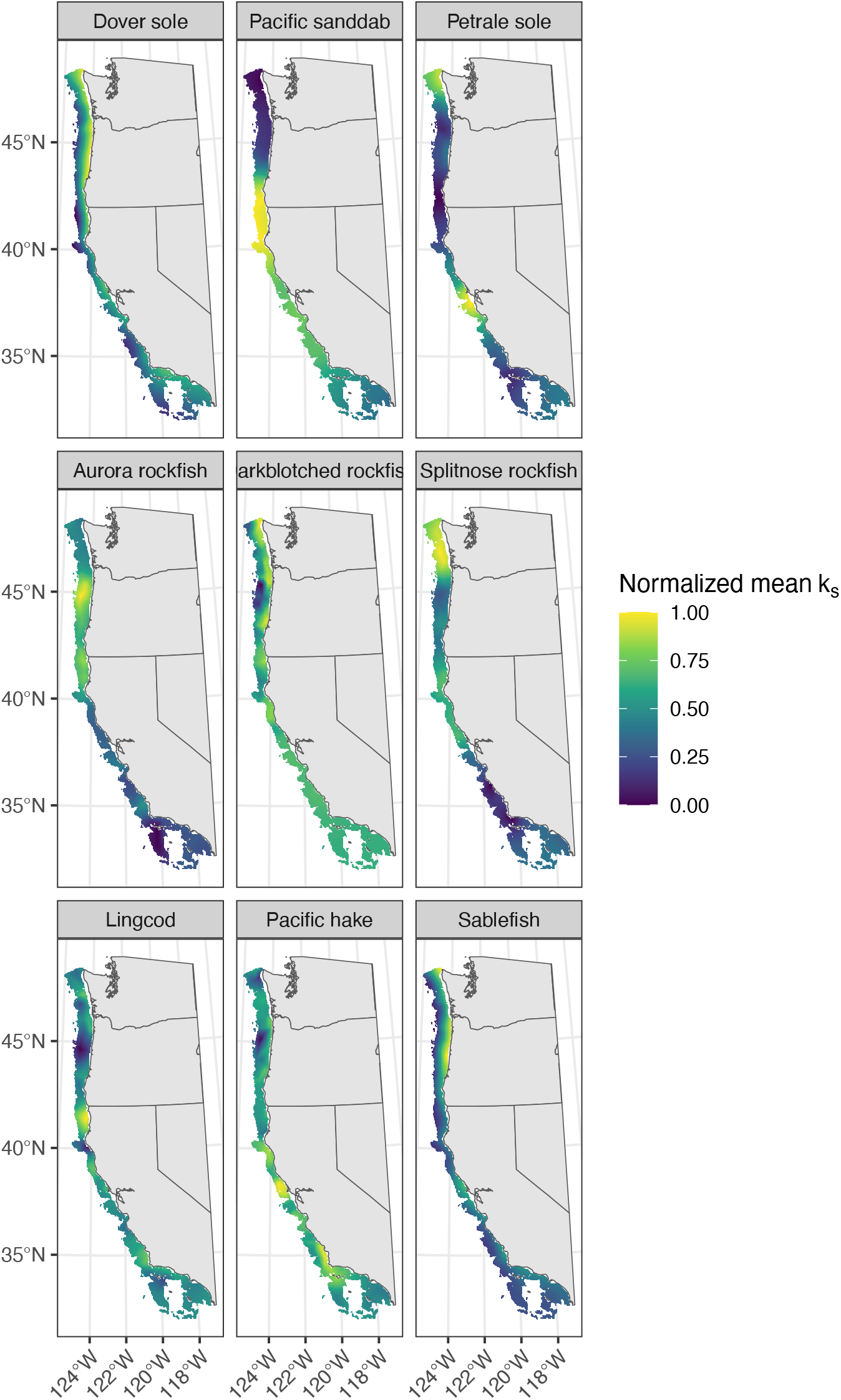
A 4×4 km gridded map of estimated growth coefficients, *k*_*s*_, for each species (panels) ordered by taxa (top row: flatfish, middle row: rockfish, bottom row: roundfish). Estimates of *k*_*s*_ were averaged across years and normalized to reflect the same scale across species, from 0 (blue; slowest estimated *k*_*s*_ for the respective species) to 1 (yellow; fastest estimated *k*_*s*_ for the respective species). The temporal coverage of data that was available varied between species.

Much like growth rate, locations with favorable body condition did not appear consistent with life history or depth distribution (Figure 4). Shelf species, which are also short-lived with early maturation, demonstrated a patchy latitudinal gradient in body condition, with individuals having decreased body condition in the northern half of their range and better body condition in the southern half, particularly in the Santa Barbara Channel. Though, more specifically, petrale sole displayed a broader spatial distribution of decreased body condition, extending from the northern end of their range to central California, while lingcod individuals with decreased body condition were primarily located off the coasts of Washington and Oregon. Slope species showed no clear patterns in body condition. Aurora rockfish displayed better body condition in the central portion of their range whereas splitnose rockfish had better body condition in the northern half. Shelf–slope species also lacked distinct patterns in body condition. Dover sole exhibited the most pronounced spatial pattern in body condition, closely mirroring its spatial pattern in growth rate (Figure S2), where individuals with decreased body condition were observed near the Washington and Oregon coastlines and individuals with better body condition were further offshore and in the southern parts of their range. Pacific hake exhibited a depth gradient, where individuals exhibited better body condition in nearshore habitats and decreased further offshore. Sablefish and darkblotched rockfish showed minimal spatial variation in body condition (Figure 2).

**Figure 4.**
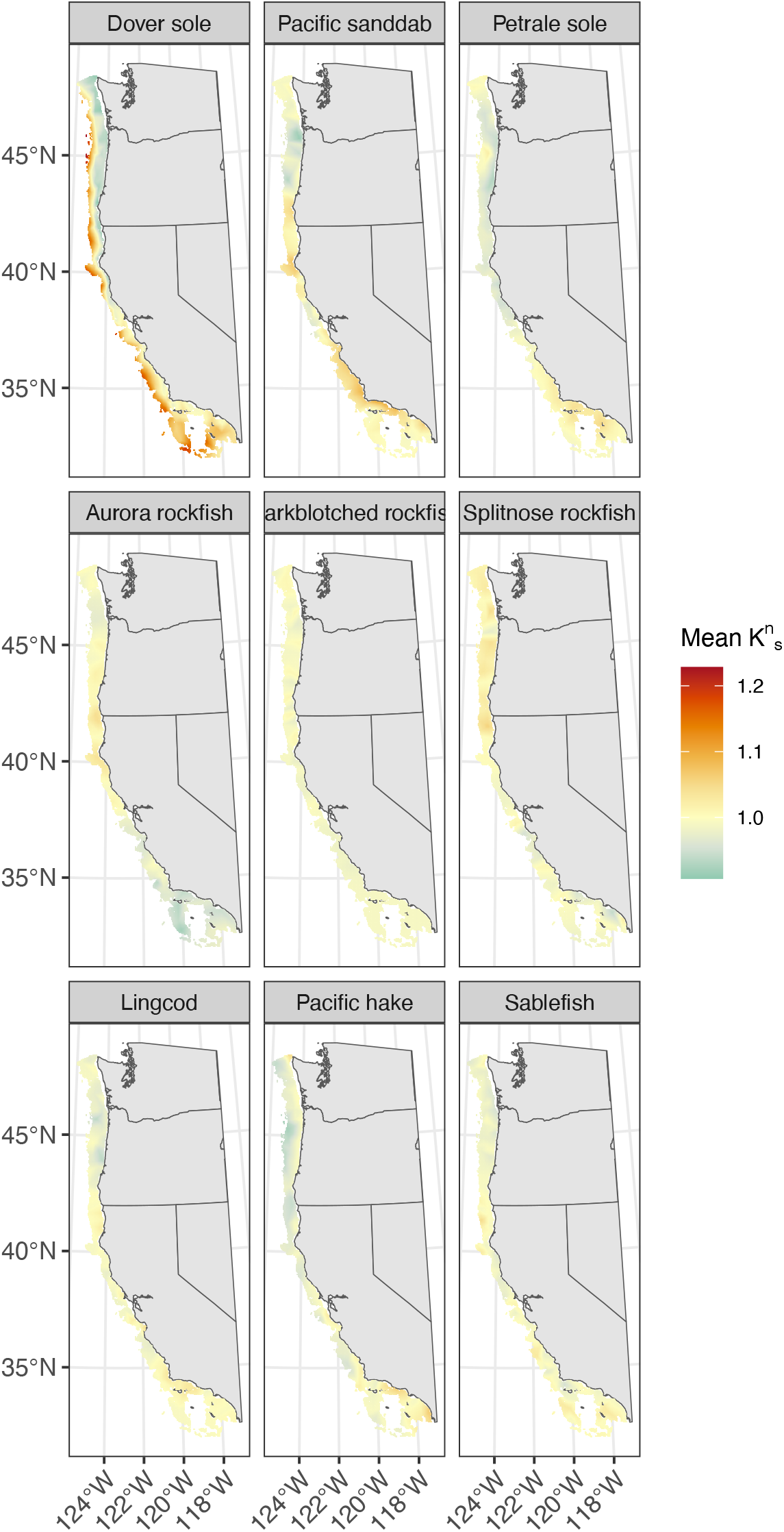
A 4×4 km gridded map of estimated body condition, 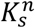, averaged across years for each species ordered by taxa (top row: flatfish, middle row: rockfish, bottom row: roundfish). Below-average body condition is indicated using blue, average body condition is indicated using yellow, and above-average body condition is indicated using red. The temporal coverage of data that was available varied between species.

### Model-Based Indices

Over time, growth rates for some species fluctuated with as little as a 10% change since 2003 (e.g., lingcod) while others fluctuated with as much as a 120% change since 2003, notably seen in darkblotched rockfish (Figure 5). The growth rates of shelf species remained at or below their historical value. Petrale sole exhibited long-term declines in their growth rate, with the sharpest decline occurring between 2015 and 2019 which coincided with a large marine heatwave. Both subpopulations of lingcod remained relatively stable around their historical value. Since 2010, the southern subpopulation of lingcod exhibited consistently higher growth rates than the northern subpopulation. Among the broadly distributed shelf–slope species, no clear pattern emerged. Pacific sanddab and sablefish initially exhibited declines in their growth rate but it eventually increased and exceeded their historical value by the end of their respective time series. In contrast, the growth rates of darkblotched rockfish, dover sole, and Pacific hake remained higher than their historical value. Growth rates of species inhabiting the slope regions remained below that of their historical value for the entirety of their time series.

**Figure 5.**
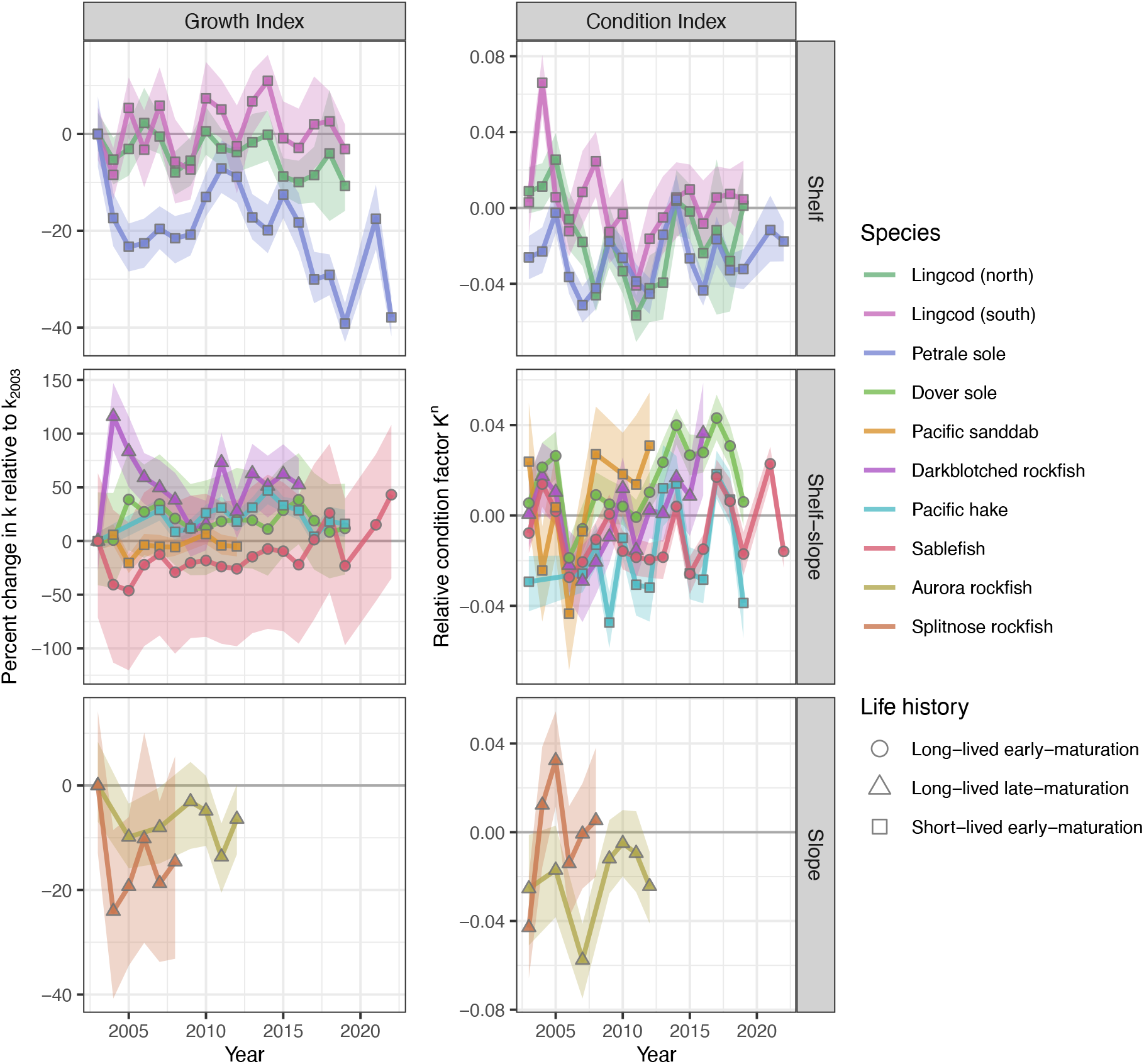
Model-based indices of growth rate *k*_*t*_ and body condition 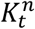. Growth rate is displayed using percent change relative to *k* in the first time step (*t* = 2003). Species (colors) are grouped by shelf (top row), shelf−slope (middle row), and slope (bottom row) species and shapes indicate life-history types. Dark-gray lines mark zero for percent change in growth rate (left column) and a value of one for body condition (right column). Shaded regions indicate 95% confidence intervals.

Body condition varied uniquely among groundfish (Figure 5). The body condition of shelf species remained either at or below average body condition since 2010. Both subpopulations of lingcod exhibited decadal fluctuations, such that their body condition declined to its lowest in 2011 followed by an improvement toward average body condition through the end of their respective time series. The body condition of the northern subpopulation of lingcod almost consistently remained below that of the southern subpopulation, likely due to latitudinal differences in productivity along the coast. Shelf–slope species tended to exhibit large interannual fluctuations with some shared peaks in body condition. All shelf–slope species appeared to experience a sharp decline in body condition in 2006 just prior to a long-term increase, except for Pacific hake in which data for that year is not available. Slope species, which are both long-lived and late maturing, also displayed large fluctuations over their short time series, though the body condition of splitnose rockfish remained higher than that of aurora rockfish for most of the time series. The other long-lived and late maturing species, darkblotched rockfish (a shelf–slope species), similarly displayed large fluctuations. All three long-lived and late maturing species exhibited sharp declines in body condition between 2005 and 2007.

## Discussion

Our analysis of growth rate and body condition across nine groundfish species showed substantial variability across space and time, indicative of the dynamic ocean environment they inhabit. Some species varied more across space while others varied more across time in a given location, but these patterns were not consistent with life history or depth distribution. We also observed no clear hotspots of favorable growth rate or body condition common across species with similar life histories or depth distributions, suggesting that local-scale responses in somatic growth might be driven more by niche partitioning. At the regional scale, species exhibited unique trends in growth rate and body condition with interspecific differences in the direction and extent of change throughout time. Together, these results suggest that these nine groundfish species are responding in largely species-specific ways despite commonalities in their life-history traits, habitats, and physical ocean conditions.

### Comparison to previous literature

In our study, we modeled variability in space continuously to better capture the true nature of ecological processes. Previous methods discretized space, often by survey stratification or management boundaries (Gertseva *et al*., 2017). However, such predetermined breakpoints might not accurately reflect the life-history structure of populations. In such cases, generalized additive models have been used to detect locations of demographic change in growth data (Kapur *et al*., 2020) but this approach does not overcome the low resolution of aggregated data. Given the region’s dynamic ocean environment, modeling spatial variation as a continuous process is likely necessary to capture the level of heterogeneity exhibited by some groundfish in the U.S. West Coast, which ultimately could improve model-based estimates and overcome limitations in sampling (Barnett *et al*., 2021; Anderson *et al*., 2024).

We observed spatial patterns in growth rate that differed from previous studies which estimated variability at coarser spatial resolution. VBGF models with spatially-varying asymptotic length (*L*_∞_) and growth rate (*k*) revealed U.S. West Coast groundfish occupying the shelf exhibited the highest growth rates between Cape Blanco and Cape Mendocino (Gertseva *et al*., 2017) likely due to favorable upwelling conditions (Parrish *et al*., 1981). While we observed similar patterns in lingcod, we found that, in contrast, petrale sole exhibited their slowest growth rate in this area. Discretizing space might have oversimplified patterns of variability for some species like petrale sole which exhibited very localized hotspots of favorable growth. However, a limitation to our study is the assumption of a static and known *L*_∞_. Evidence suggests that *L*_∞_ tends to decline with warming temperatures (Gertseva *et al*., 2017; Ikpewe *et al*., 2021) and ignoring such information can affect estimates of *k*and further contribute to differences in spatial patterns of growth rate between our studies.

Our findings of spatiotemporal variability in body condition parallel that of an extensive analysis conducted for 28 groundfish species (Thorson, 2015). They found that a substantial fraction of variation can be attributed to annually-varying spatial differences (i.e. spatiotemporal), suggesting that variability is driven largely by local environmental conditions. In our study, we find similar hotspots of favorable body condition, most notably for dover sole, despite our exclusion of environmental covariates such as temperature and season. The magnitude of the environmental covariates they included, however, was small relative to that of spatial and spatiotemporal variation, suggesting that other unobserved variables are influencing the body condition of groundfish.

### Biological and management insights

We find that niche partitioning might play a large role in the variability of somatic growth in groundfish along the U.S. West Coast, obscuring the anticipated effect of life history and depth distribution. A combination of environmental, ecological, and fishing factors can drive local habitat conditions in which many species of groundfish coexist. One mechanism for coexistence between sympatric species with similar life-history traits is resource partitioning along one or more niche dimensions (Macarthur and Levins, 1967), most notably in food resources for marine fish (Ross, 1986). Trophic partitioning in a given location can depend on factors such as prey availability (Brodeur and Pearcy, 1986), size structure (Barnes *et al*., 2021), and dietary preferences (Knickle and Rose, 2014). Variation in these factors across space and time are likely to drive fine-scale variability in somatic growth between similar species.

Regional-scale annual indices of somatic growth metrics were also generally unique among species, with few linkages to certain life-history traits and depth distributions. Shared peaks and dips in the annual indices of body condition were observed for species within similar depth distributions which suggests regional-scale climate conditions might influence directional change in body condition of species at similar depths regardless of their life-history traits. In contrast, for annual indices of growth rate, few similarities were observed across species, which supports previous findings that regional-scale climate conditions might not be its primary driver (Stawitz *et al*., 2015). However, because some species might respond quickly to environmental change, it would be valuable to explore trends on shorter timescales to fully discern whether similar species have shared responses to environmental change.

A statistical evaluation of consistent patterns among species with similar life history or depth distribution was limited by the availability and sample size of measurements of individual fish. Historically, sampling effort has varied between species. As these data are primarily used in stock assessments, species with recent assessments have more recently collected data to inform their growth estimates. Limited sampling overlap across all nine groundfish species in years with extreme environmental conditions (e.g., marine heatwaves in 2014, 2019) make it difficult to evaluate support for consistent responses to such events among similar species. Consistent monitoring across years and species or targeted monitoring during years with extreme environmental conditions may improve the capacity to evaluate drivers of growth variability and forecast growth under environmental change.

We found that short-lived species were among those that exhibited relatively low levels of variability in body condition. However, short-lived species are usually expected to track changes in the environment more closely than long-lived species, and thus, we would expect that their body condition fluctuates relatively more than long-lived species (Bjørkvoll *et al*., 2012). They have a greater capacity to compensate for environmental variability through changes in their growth which allows them to quickly stabilize following environmental change, a process known as compensatory growth (Ali *et al*., 2003). When compensatory growth is rapid, using size measurements sampled annually might not capture the short timescale in which compensatory responses are occurring. Consequently, it is possible that accelerated growth typically exhibited by short-lived species following periods of growth depression might appear stable when observed at coarser timescales because of their ability to quickly re-stabilize. Directly measuring growth of individuals over days or weeks might reveal greater fluctuations in short-lived species compared to long-lived species, aligning more closely with our expectation that short-lived species have greater plasticity and fluctuations in their growth. However, many commercially harvested groundfish species are a challenge to maintain in the lab settings necessary to conduct such studies.

Consideration of variability in growth within stock assessments for fisheries management is receiving greater recognition (Lorenzen, 2016; Maunder *et al*., 2016; Andersen, 2020; Punt *et al*., 2020) and our findings contribute to the understanding of potential mechanisms driving such variability in groundfish. Within stock assessments, species are often managed as a single unit across a large spatial domain and it is assumed that growth is constant, which can lead to inaccurate and biased estimates of spawning biomass, recruitment, and spawning depletion (Kerr *et al*., 2017; Stawitz *et al*., 2019). Incorporating variability in growth has been shown to minimize inaccuracies (Correa *et al*., 2021) through improved estimates of age compositions (Correa *et al*., 2020). Therefore, leveraging information regarding variability in growth can improve stock assessment estimates through methods such as the empirical weight-at-age approach and temporally- or spatially-varying parameters in growth models.

Our findings could inform the inclusion of variability in growth into stock assessments in a variety of ways. Species with clear spatial differences in growth rate can be discretized into subpopulations with unique parameters estimated for each to reflect its spatial variability. Of the species investigated, the stock assessment for lingcod is currently the only assessment that accounts for genetic differences by splitting the population into northern and southern subpopulations (Haltuch *et al*., 2018). However, stock assessments for other species with clear spatial differences in growth rate, such as the latitudinal gradient exhibited in Pacific sanddab and petrale sole, might also benefit from estimating unique parameters within subpopulations. Identifying potential environmental covariates that best explain the observed variability in growth to inform forecasts that include environmental variability (Punt *et al*., 2021) may be more important than discretizing populations into subpopulations. Careful consideration of environmental covariates and rigorous testing of its performance can guide its integration into stock assessments. The importance of integrating variability in growth into stock assessments will depend on its influence on assessment estimates which might differ among species (Lorenzen, 2016).

Our findings might also facilitate the prioritization of certain locations for data collection. We did not observe consistent patterns of variability in growth among species with similar life history or depth distributions, which suggests that data collection efforts from one species or region might not be useful to extrapolate to other species or regions. Results from the spatiotemporal models used here can reveal where uncertainty or variability are the highest and those locations might deserve more attention with respect to sampling. Historically, the locations of samples within a survey had to be randomly selected to maintain the statistical validity of results but clustered or concentrated sampling can now be accommodated with spatiotemporal models (Thorson *et al*., 2020; Vilas *et al*., 2024). Data collected could reflect the breadth of variability such that areas with high variability would be sampled more heavily than areas that vary minimally. However, sampling more at specified locations might only be useful for externally estimating growth, as it will provide more informative estimates without impacting those of selectivity. Generally, when thinking about changing the design of a survey, given inevitable tradeoffs, maintaining some level of randomly located samples can capture the array of environmental and demographic characteristics that influence the variation in populations’ productivity across space.

## Conclusion

Variability in somatic growth is ubiquitous across the groundfish species explored here, but weak associations to life history and depth distributions challenge the idea that similar species will exhibit shared responses to broadly experienced environmental conditions. Exploring the effect of niche-partitioning on growth with the use of ecosystem models is a clear next step toward understanding the mechanisms driving variability in growth. At the time of analysis, some species had limited age information in recent years due to backlogs in sample processing. Maintaining up-to-date biological sampling and aging will increase the richness of available data for these species and result in a more comprehensive time series for the development of model-based indices of somatic growth. The dynamic nature of somatic growth suggests that current assumptions of its uniformity might bias estimates of population status, and consideration of its variability is likely to improve estimates of productivity for more robust and climate-ready fisheries.

## Acknowledgements

The authors would like to thank the Northwest Fishery Science Center for providing the data for this analysis as well as Megan L. Feddern, Owen Hamel, and an anonymous reviewer for their helpful feedback.

## Data Availability

The data underlying this article is available in GitHub at https://github.com/AndreaNOdell/groundfish_growth

## Author Contributions

Study conception by A.N.O and E.J.W. Formal analysis conducted by A.N.O. with input from K.N.M, E.J.W, and K.F.J. Original draft of manuscript prepared by A.N.O with revisions and edits provided by all authors.

## Competing Interests

The authors declare no competing interests.

## Funding

Funding for this project came from a NMFS-Sea Grant Population and Ecosystem Dynamics Fellowship (award #NA23OAR4170535) to A.N.O, K.N.M, and M.L.B. and a National Science Foundation National Research Traineeship (award #1734999) to A.N.O. M.L.B acknowledges support from University of California Agricultural Experiment Station (Project #CA-D-ESP-2244-H).

## Notes

### Competing Interest Statement

The authors have declared no competing interest.

