## Supplementary Materials for "Spatiotemporal Models Reveal Dynamic Growth Patterns in U.S. West Coast Groundfish"

### Distribution of samples across species

We obtained 54,148 samples total. Sample sizes for each species were spread evenly across available years in most cases (Table S1). Total annual sample size fluctuated due to lack of samples for some species in some years (Table S1). Lack of samples in a given year was often attributed to delays in assessing otoliths for age information. Ageing via otoliths is commonly conducted in preparation for a stock assessment, so species without recent assessments typically have fewer aged otoliths and consequently a truncated timeseries. Total sample size for each species were greater than 1234 samples. More abundant species were associated with larger sample sizes. The age distribution of samples varied among species depending on lifespan. Short-lived species were well-sampled across age classes with fewer samples of the youngest and oldest age classes as expected. Species with longer lifespans had weaker representation in the oldest age classes, resulting in longer tails and right-skewness (Figure S3). Species were sampled throughout their geographic range which primarily ranged from the Canada-US border to the Mexico-US border (Figure S4). Rarer species, such as rockfish, had sparser spatial resolution compared to more abundant species such as sablefish, and darkblotched rockfish are not typically observed south of Point Conception, California.

### Estimating $\bar{w}$

To estimate condition factor $K_n$, we first needed to estimate the expected weight $\bar{w}$ for a given length $l$ across the entire survey period in the absence of spatial or temporal variability. We modeled $\bar{w}$ as

1. $log(\overline{w}) \sim Student(\mu, \phi, \nu)$,
2. $\mu = x + y\, l$,

where $\mu$ represents the mean weight, $\phi$ represents the scale parameter, $\nu$ represents the available degrees of freedom which we set to 5 (Lindmark *et al.*, 2022a). For each species, we estimated an intercept $x$ and slope $y$ using log-transformed lengths $l$.

### Supplementary Figures

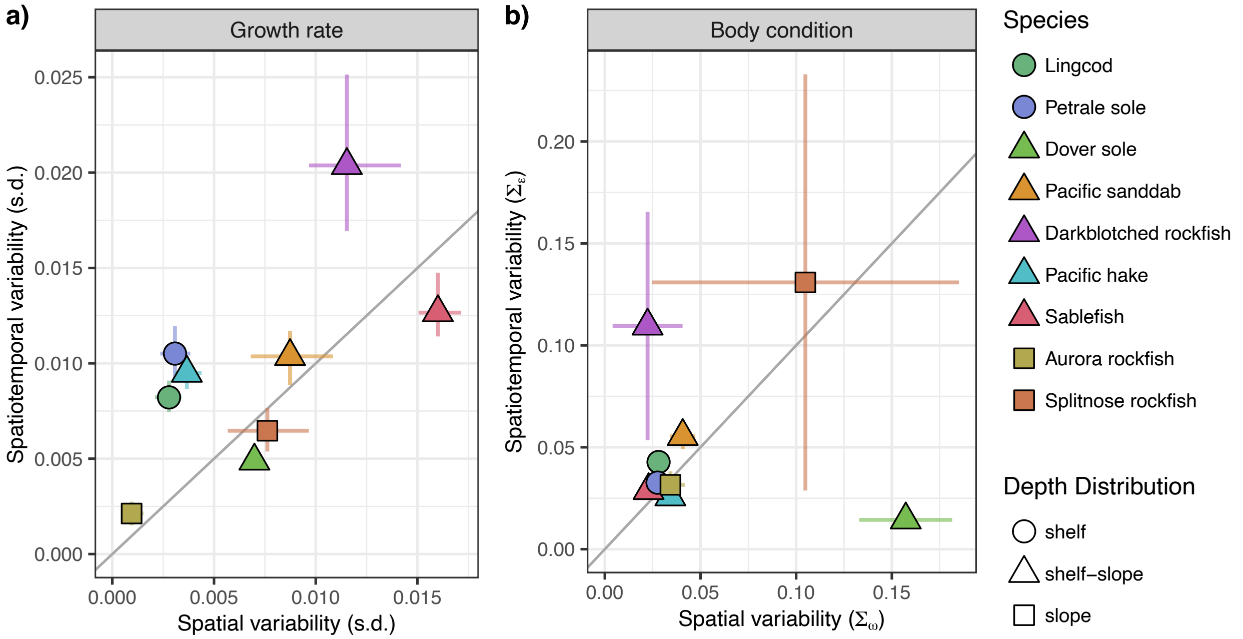


Figure S1. The range of spatial and spatiotemporal variability for growth rate *k* and body condition $K^n$ for species (colors) that inhabit the shelf (circles), shelf and slope (triangles), or the slope (squares). Variability in $k$ is measured as the standard deviation (s.d.) calculated from estimates of *k* and variability in $K^n$ is measured as the standard deviation of the spatial and spatiotemporal fields estimated by the body condition model,

$\Sigma_\omega$ and $\Sigma_\varepsilon$, respectively. The 1:1 line indicates whether spatiotemporal variability (above) or spatial variability (below) is more influential.

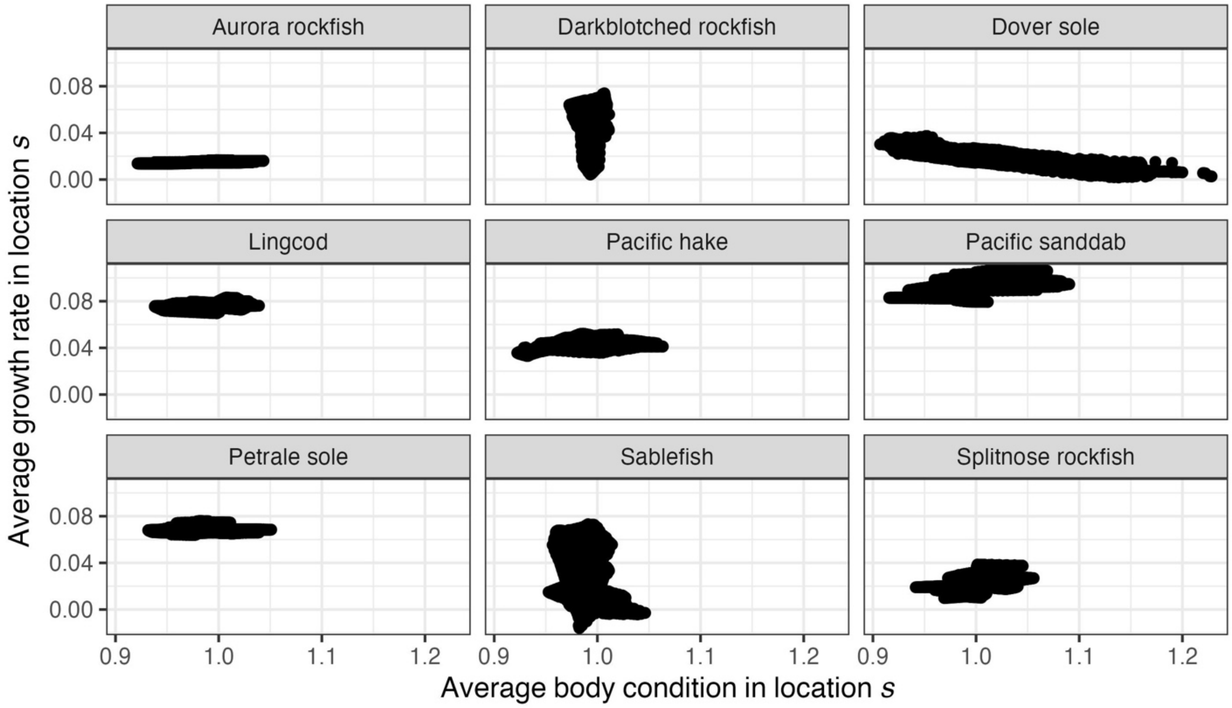


Figure S2. The relationship between the average growth rate and average body condition in each location. Each point represents one 4x4 km location.

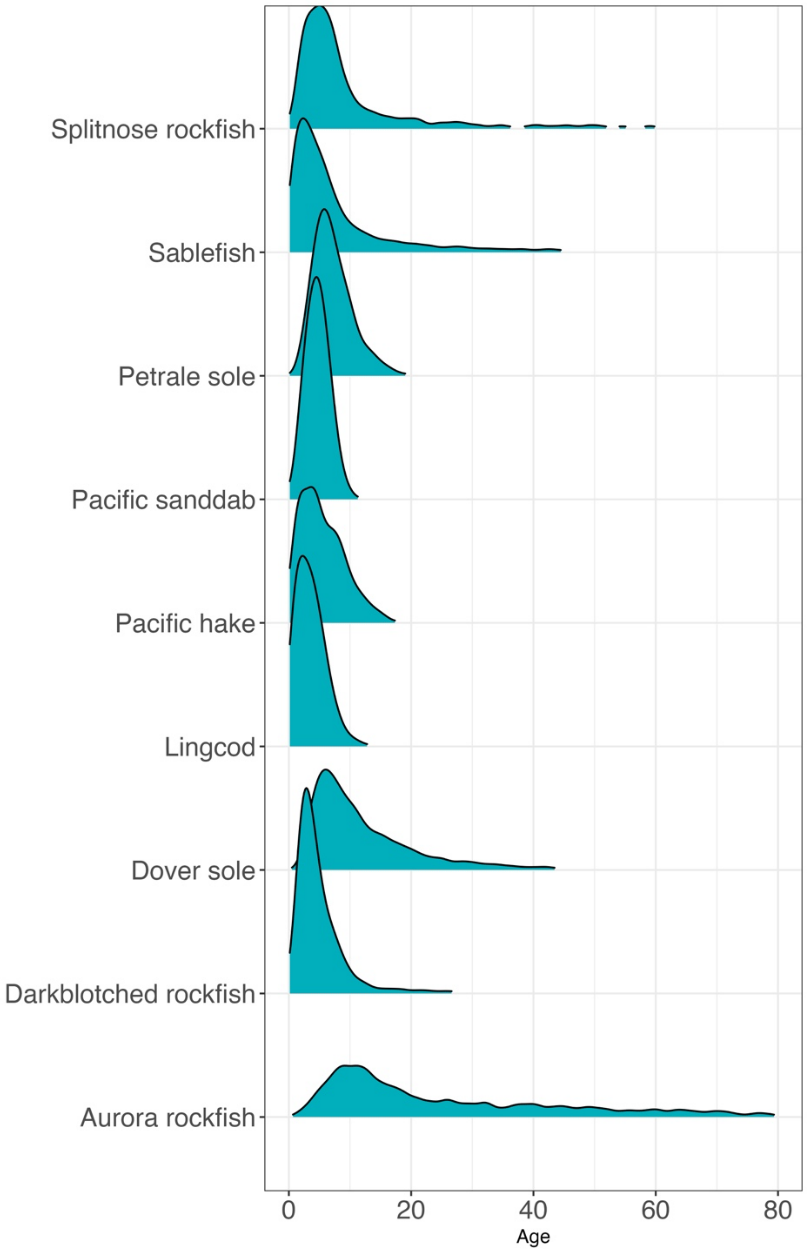


Figure S3. Distribution of available data across age classes for each species

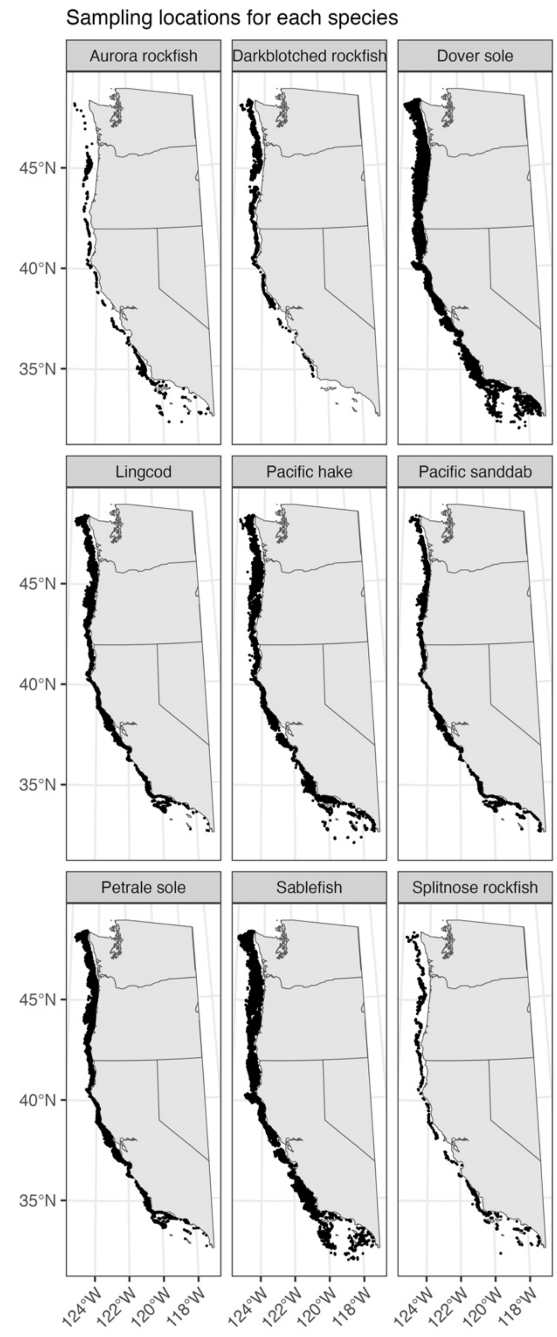


Figure S4. Map of sampling locations aggregated across years for each species. Sparser spatial sampling for some rockfish species is associated with their relative rarity compared to other more abundant species. In general, latitudinal range is comparable across species, except for darkblotched rockfish which rarely occupy habitat south of Point Conception, California.

### Supplementary Tables

Table S1. Total annual sample size of paired length, weight, and age measurements for females of each species. Empty cells indicate years in which zero paired length, weight, and age measurements for females are available.

| | **2003** | **2004** | **2005** | **2006** | **2007** | **2008** | **2009** | **2010** | **2011** | **2012** |
|---|---|---|---|---|---|---|---|---|---|---|
| **aurora rockfish** | 222 | | 207 | | 206 | | 200 | 226 | 236 | 252 |
| **darkblotched rockfish** | 330 | 264 | 357 | 466 | 468 | 350 | 519 | 439 | 385 | 377 |
| **dover sole** | 522 | 446 | 504 | 518 | 562 | 540 | 588 | 589 | 603 | 634 |
| **lingcod** | 535 | 463 | 505 | 468 | 330 | 430 | 258 | 304 | 311 | 238 |
| **Pacific hake** | 896 | | | | 777 | 592 | 358 | 373 | 335 | 330 |
| **Pacific sanddab** | 501 | 957 | 628 | 462 | 525 | 518 | | 640 | 476 | 538 |
| **petrale sole** | 382 | 430 | 375 | 419 | 381 | 398 | 399 | 396 | 398 | 398 |
| **sablefish** | 681 | 519 | 732 | 612 | 580 | 490 | 508 | 520 | 569 | 529 |
| **splitnose rockfish** | 227 | 192 | 217 | 197 | 197 | 204 | | | | |
| **Year total** | 4296 | 3271 | 3525 | 3142 | 4026 | 3522 | 2830 | 3487 | 3313 | 3296 |

| | **2013** | **2014** | **2015** | **2016** | **2017** | **2018** | **2019** | **2021** | **2022** | **Species total** |
|---|---|---|---|---|---|---|---|---|---|---|
| **aurora rockfish** | | | | | | | | | | 1549 |
| **darkblotched rockfish** | 326 | 330 | 475 | 336 | | | | | | 5422 |
| **dover sole** | 458 | 648 | 644 | 643 | 653 | 651 | 274 | | | 9477 |
| **lingcod** | 251 | 278 | 278 | 219 | 156 | 267 | 178 | | | 5469 |
| **Pacific hake** | 218 | 345 | 399 | 452 | 492 | 423 | 173 | | | 6163 |
| **Pacific sanddab** | | | | | | | | | | 5245 |
| **petrale sole** | 477 | 400 | 398 | 501 | 523 | 476 | 367 | 467 | 504 | 8089 |
| **sablefish** | 452 | 566 | 586 | 582 | 588 | 730 | 415 | 1042 | 798 | 11499 |
| **splitnose rockfish** | | | | | | | | | | 1234 |
| **Year total** | 2182 | 2567 | 2780 | 2733 | 2412 | 2547 | 1407 | 1509 | 1302 | 54,147 |

Table S2. Estimates of maximum age and age at 50% maturity for each species.

| **Species** | **Maximum Age** | **Age at 50% Maturity** | **Reference** |
|---|---|---|---|
| Aurora rockfish | 125 years | 13 years | Hamel *et al.,* 2013 |
| Dover sole | 69 years | 5 years | Wetzel and Berger, 2021 |
| Pacific sanddab | 13 years | 3 years | He *et al.,* 2013 |
| Petrale sole | 31 years | 5 years | Taylor *et al.,* 2023 |
| Darkblotched rockfish | 98 years | 8 years | Wallace and Gertseva, 2017 |
| Splitnose rockfish | 103 years | 9 years | Gertseva *et al.,* 2009 |
| Lingcod | 13 years | 3 years | Taylor *et al.,* 2021 |
| Pacific hake | 25 years | 4 years | Berger *et al.,* 2023 |
| Sablefish | 102 years | 5 years | Johnson *et al.,* 2023 |
